# Sickness engrams modulate anticipatory immune responses

**DOI:** 10.1101/2025.11.03.686140

**Authors:** Aaron Douglas, Andrea Muñoz Zamora, Paul Conway, Esteban Urrieta, Lydia Lynch, Tomás J Ryan

## Abstract

A threat to survival in the wild is vulnerability to infection. The immune system is essential for defence against foreign species which cause sickness. During infection the brain triggers conserved behaviors, including fever, tiredness and anorexia that support immunity. The immune system stores infection information via adaptive immunity, however it remains unclear whether the brain stores immune-related information as long-term memory engrams. Here we demonstrate that mice form contextual memories for sickness events. Upon sickness-memory recall mice lower whole-body metabolism, and increase coactivation between the hippocampus and sickness regions such as the central amygdala, alongside elevated engram activation. Optogenetic reactivation of sickness engrams decreases metabolism, similar to natural recall. Finally, natural recall and artificial reactivation of a sickness-memory increased genes associated with the acute phase response in the liver. These findings suggest that sickness experiences are encoded as engrams, which upon reactivation trigger coordinated metabolic and innate immune responses.

## Introduction

Brain-body interactions are fundamental to organismal physiology, with the brain coordinating peripheral signals to maintain homeostasis^1,2^. Mounting research highlights the bidirectional nature of brain-body communication, such as mental state influencing recovery from cancer and myocardial infarction, while peripheral signals modulate the circadian rhythm of sleep and anxiety behaviours^3–6^. A major aspect of brain-body interactions is the crosstalk between the nervous and immune systems^7^. Neuroimmune interactions convey information about peripheral infection and inflammation to the brain. For example, cytokines released from immune cells modulate the excitability of peripheral neurons, particularly in the nodose ganglion, informing the brain about the inflammatory environment^8^. Neurons in the nucleus of the solitary tract (NST) respond to peripheral inflammation by modulating the cytokine response, a mechanism known as the inflammatory reflex^9,10^. Furthermore, cytokines such as interleukin-1beta (IL-1β) enhance the excitability of neurons in the ventromedial preoptic area (VMPO) directly, to generate fever during sickness^11^. Together, these studies describe an essential role for neuroimmune crosstalk to generate an adequate response to inflammation and infection. However, whether the brain can store long-term information about sickness or immune events for later recall remains unclear.

Previous studies have shown that specific ensembles of neurons are reactivated during subsequent inflammatory events^12^. In particular, ensembles within the agranular insular cortex (aIC) encode distinct information about inflammatory diseases such as colitis and peritonitis. Reactivation of these ensembles triggers innate immune responses tailored to each condition^12,13^. These findings raise the possibility that the brain may not only store disease-specific immune representations, but also use this information to proactively prepare for anticipated threats, described as anticipatory immunity. However, it is unknown whether such ensembles reflect innate signatures for immune responses, or whether the brain can form “immune engrams” though learning, for memory-guided, predictive immune activation ^8,10,11^ The concept of anticipatory immunity originated in 1885 with the rose-cold case study, which described a woman exhibiting an allergic reaction to an artificial flower^14,15^. Since then, anticipatory immunity has been demonstrated to exist across multiple species, female bed bugs upregulate immunity in anticipation of copulation; echinoderms increase circulating immune cell density in response to perceived predation risk; and fish double their circulating lymphocyte counts when exposed to perceived threats^16–18^. Numerous studies have aimed to classically condition the immune system by pairing an immunological unconditioned stimulus (US) and a neutral conditioned stimulus (CS)^19–21^. A neural basis for immune conditioning has been established by a recent study that paired sweetened water (CS) with the bacterial endotoxin lipopolysaccharide (LPS, US), inducing conditioned taste aversion through a signalling axis in the aIC^22^. However, it remains unclear whether such responses rely on memory engram reactivation, and whether the immune system is pre-emptively activated.

Here, we investigated whether mice form contextual memories for sickness events in the form of engrams. Engrams can be defined as neuronal ensembles that are activated by learning and undergo physical changes, and the reactivation of which leads to retrieval of the target memory^23–27^. We demonstrate that mice learn to associate a particular novel context with a sickness experience induced by the bacterial compound LPS. Re-exposure to the sickness context led to a reduction in whole-body metabolism and increased expression of acute phase response proteins (APPs) in the liver. Furthermore, during sickness memory recall, hippocampal regions show increased coactivation with classical sickness-related areas, including the central amygdala (CeA), paraventricular nucleus of the thalamus (PVT), and paraventricular nucleus of the hypothalamus (PVH). Additionally, we show that sickness engrams form in the hippocampal dentate gyrus (DG), and in sickness associated regions. Finally, artificial reactivation of DG sickness engrams is sufficient to elicit a decrease in whole-body metabolism and an increase in liver APPs, similar to the physiological response to LPS. Together, these findings suggest that hippocampal sickness engrams are sufficient to drive both metabolic and anticipatory immune responses.

## Results

### Contextual memory recall of a sickness event decreases whole-body metabolism

Sickness behaviours are an evolutionarily conserved set of actions that include fever, inactivity, social withdrawal and decreased food consumption^28^. Additionally, inflammatory compounds decrease whole-body metabolism as an energetic trade-off to fuel the immune response against infection^29^. Although sickness behaviours occur across different inflammatory molecules and infections, the degree to which metabolism and behaviour are affected varies depending on the specific microbe or compound^29^. The toll-like receptor four (TLR4) activator LPS is a potent modulator of both sickness behaviours and whole-body metabolism compared to other TLR agonists^29^. Therefore, we used the behavioral and metabolic effects of LPS as measurable readouts to assess sickness memory formation.

To investigate whether mice can remember a contextual area in which they have previously been sick, we performed a conditioned place avoidance paradigm (CPA, Fig. 1a). Mice were injected with LPS and confined to one side of the apparatus for 4 hours, followed by a 2-day recovery period, which is sufficient for full recovery from this dose^30^. Mice previously injected with LPS avoided the side of the CPA arena where they had been sick, while PBS injected mice showed no preference (Extended Data Fig. 1a; Fig. 1b,c), as previously described^31^.

**Figure 1:**
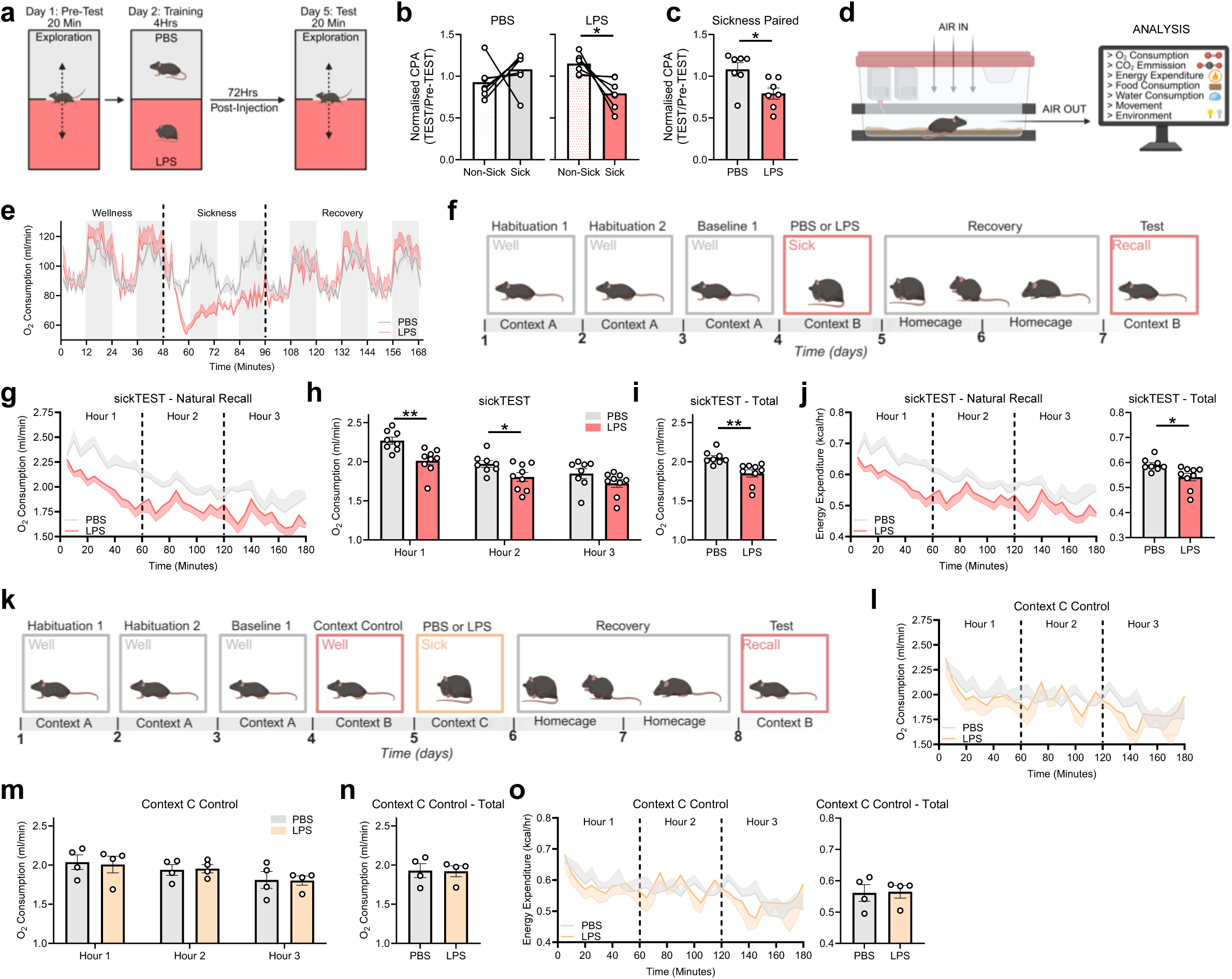
A sickness-associated contextual memory modulates whole-body metabolism. **a**, Experimental timeline of CPA paradigm. **b**, Normalised place preference for PBS-conditioned aversion (left) and LPS-conditioned aversion (right; Test/Pre-test). **c**, Normalised place preference for PBS-paired side (grey) versus LPS-paired side (red). **d**, Schematic of metabolic cage analysis. **e**, Time plot of oxygen consumption between PBS-injected mice (grey) and LPS injected mice (red). **f**, Experimental timeline of sickness training paradigm. **g**, Time plot of oxygen consumption between PBS-injected mice (grey) and LPS-injected mice (red) on sickness recall day. **h**, Comparison of oxygen consumption between PBS-injected mice (grey) and LPS-injected mice (red) on sickness recall day at hour 1, 2 and 3, with **i**, total time averaged in metabolic cages. **j**, Time plot of energy expenditure between PBS-injected mice (grey) and LPS-injected mice (red) on sickness recall day (left), with total time averaged in metabolic cages (right). **k**, Experimental timeline of context C control training paradigm. **l**, Time plot of oxygen consumption between PBS-injected mice (grey) and LPS-injected mice (orange) when replaced into context B. **m**, Comparison of oxygen consumption between PBS-injected mice (grey) and LPS-injected mice (orange) when replaced into context B at hour 1, 2 and 3, with **n**, total time averaged in metabolic cages. **o**, Time plot of energy expenditure between PBS-injected mice (grey) and LPS-injected mice (orange) on sickness recall day (left), with total time averaged in metabolic cages (right). Data are shown as mean ± s.e.m, n = 4–9 mice per group. Significance was calculated using a two-tailed paired students t-test (**b)**, or a two-tailed unpaired students t-test (**c,h-j,m-o**). \**P* < 0.05, \*\**P* < 0.01.

To determine whether a sickness memory is sufficient to decrease whole-body metabolism, we used metabolic cage technology to measure oxygen consumption, energy expenditure, food and water consumption, and activity (Fig. 1d). Consistent with previous reports, LPS administration decreased oxygen consumption (Fig. 1e), carbon dioxide emission (Extended Data Fig. 1b), energy expenditure (Extended Data Fig. 1c), food consumption (Extended Data Fig. 1d), water consumption (Extended Data Fig. 1e) and movement (Extended Data Fig. 1f,g)^29^. We next developed a novel Pavlovian conditioning timeline, in which mice were trained to associate context A with wellness and context B with either LPS-induced sickness or a saline injection (Fig. 1f). Upon re-exposure to context B after recovery, mice that had received LPS exhibited a significant reduction in oxygen consumption compared to PBS controls (Fig. 1g). This reduction persisted for 2-hours, and remained lower over the total period of 3-hours (Fig. 1h,i). Similarly, LPS-conditioned mice also showed decreased energy expenditure (Fig. 1j) and carbon dioxide emission (Extended Data Fig. 2a), along with an unexpected increase in food consumption (Extended Data Fig. 2c). Together, these results suggest that mice form a memory of prior sickness, which can be recalled to alter behaviour and whole-body metabolism.

To confirm the decrease in whole-body metabolism was specific to the sickness-associated context and not due to prolonged metabolic effects of LPS, we introduced a third context into our Pavlovian conditioning timeline (Fig. 1k). Mice were trained to associate context A and B with wellness, and received LPS or PBS injections paired with context C. Upon re-exposure to context B, we observed no difference in oxygen consumption between LPS-treated and PBS control mice (Fig. 1l-n), as sickness was paired with context C. Similarly, no differences were observed in energy expenditure (Fig. 1o), carbon dioxide emission (Extended Data Fig. 2a) or food consumption (Extended Data Fig. 2c). Furthermore, respiratory exchange ratio and activity levels did not differ between mice conditioned with context B or context C, suggesting that these readouts are not context dependent (Extended Data Fig. 2b,c). These data demonstrate that the decrease in whole-body metabolism observed during memory recall is context specific and not a general long-term effect of LPS.

### Contextual sickness memory recall modulates liver APPs

Infection or exposure to inflammatory molecules initiates a systemic reaction known as the acute phase response^32^. The acute phase response is mediated by TLR activation on specialized immune cells and leads to the release of proinflammatory cytokines such as IL-1β, IL-6 and tumour necrosis factor (TNF)^33^. Among these, IL-6 modulates the expression of specific proteins in the liver, referred to as acute phase response proteins (APPs). These changes result in elevated levels of positive APPs (C-reactive protein; CRP) and reduced levels of negative APPs (albumin) in the serum (Fig. 2a), ultimately acting to resolve inflammation^34^.

**Figure 2:**
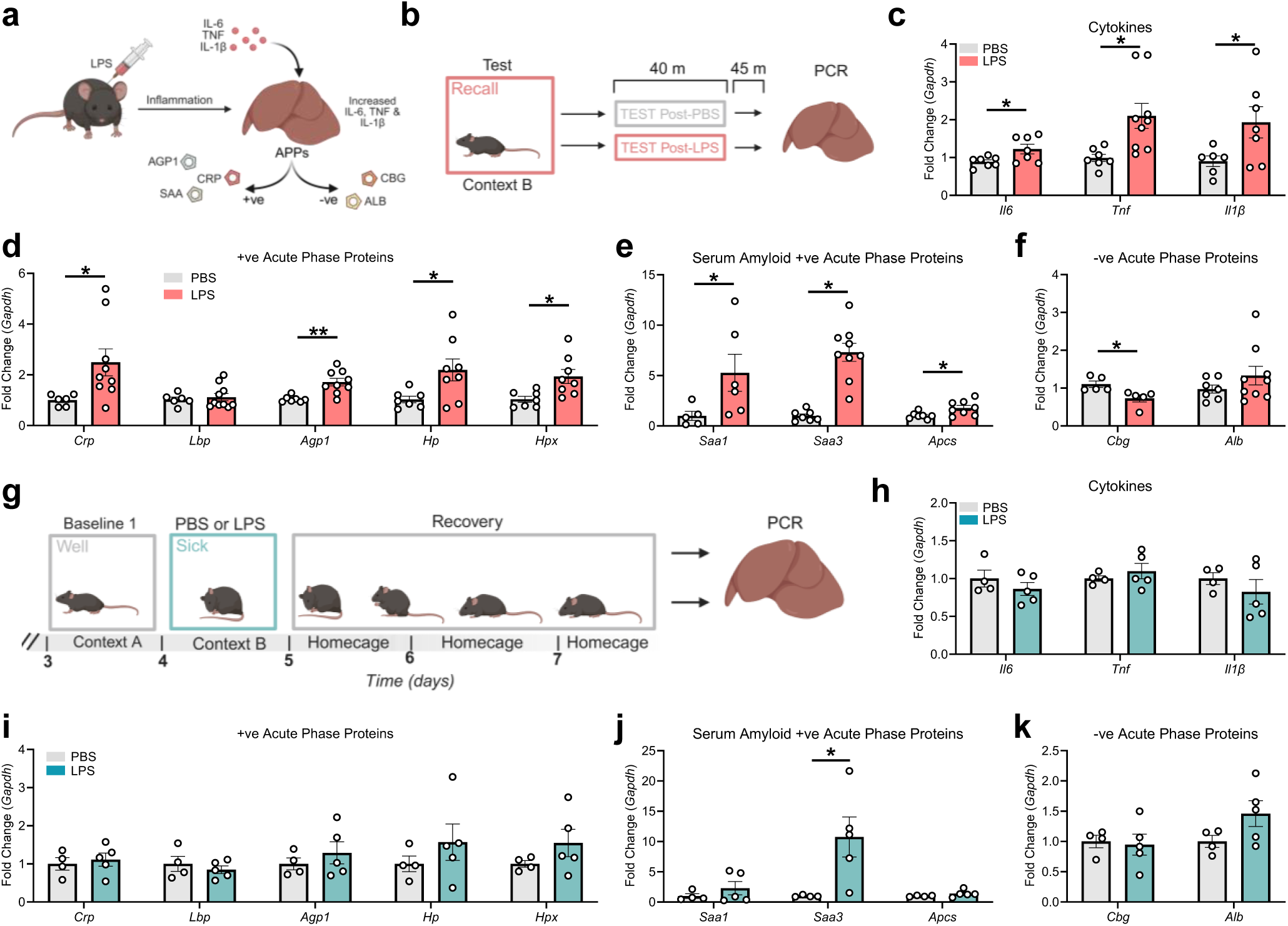
A sickness-associated contextual memory modulates acute phase response proteins in the liver. **a**, Schematic of the acute phase response to LPS in the liver. **b**, Experimental timeline for tissue collection on test day. **c**, Relative expression of the cytokines *Il6*, *Tnf* and *Il1β* in the liver of PBS-(grey) or LPS-injected (red) mice on recall day. Relative expression of the positive APPs *Crp*, *Lbp*, *Agp1*, *Hp* and *Hpx* (**d**), the serum amyloid positive APPs *Saa1*, *Saa3*, and *Apcs* (**e**), and the negative APPs *Cbg* and *Alb* (**f**) in the liver of PBS-(grey) or LPS-injected (red) mice on recall day. **g**, Experimental timeline for prolonged effects of LPS. **h**, Relative expression of the cytokines *Il6*, *Tnf* and *Il1β* in the liver of PBS-(grey) or LPS-injected (green) mice 3 days post-LPS injection. Relative expression of the positive APPs *Crp*, *Lbp*, *Agp1*, *Hp* and *Hpx* (**i**), the serum amyloid positive APPs *Saa1*, *Saa3*, and *Apcs* (**j**), and the negative APPs *Cbg* and *Alb* (**k**) in the liver of PBS-(grey) or LPS-injected (green) mice 3 days post-LPS injection. Data are shown as mean ± s.e.m, n = 4–10 mice per group. Significance was calculated using a two-tailed unpaired students t-test (**c-f,h-k**). \**P* < 0.05, \*\**P* < 0.01.

As a sickness memory could decrease whole-body metabolism similar to real-time illness, we next investigated whether it could also modulate hepatic APPs in anticipation of sickness. As expected, LPS administration induced a strong acute transcriptional upregulation of the cytokines *Il1β, Il6* and *Tnf* compared to PBS controls, measured 3 hours post-injection (Extended Data Fig. 3a). This was accompanied by robust increases in positive APPs, including *Crp*, serum amyloid A1 and A3 (*Saa1* and *Saa3*), serum amyloid P component (*Apcs*), lipopolysaccharide binding protein (*Lbp*), alpha-1-acid glycoprotein (*Agp1*), haptoglobin (*Hp*) and hemopexin (*Hpx*), along with a decrease in negative APPs albumin (*Alb*) and corticosteroid binding globulin (*Cbg*; Extended Data Fig. 3b,c).

To determine whether sickness memory alone could alter hepatic APP expression, we exposed mice to the behavioural timeline, and harvested liver tissue 45 minutes after the peak difference in oxygen consumption between LPS- and PBS-trained mice on test day (Fig. 2b). Remarkably, recall of a contextual sickness memory was sufficient to significantly increase expression of *Il6*, *Il1β*, *Tnf*, *Crp*, *Saa3*, *Apcs*, *Agp1*, *Hp*, and *Hpx*, and decrease *Cbg* expression, specifically in LPS-conditioned mice (Fig. 2c-f). To control for potential residual effects of LPS, a separate cohort underwent the same behavioural training but was not re-exposed to context B for recall (Fig. 2g). We observed no differences in APP expression between LPS- and PBS-conditioned mice, except for *Saa3* (Fig. 2h-k). These findings demonstrate that recall of a contextual sickness memory is sufficient to modulate hepatic cytokine and APP expression in an anticipatory manner.

### Sickness memories increase coactivation between the hippocampus and sickness regions

We next examined which brain regions might play a role in sickness memory encoding and retrieval. We used c-Fos as a marker for neuronal activation, and compared c-Fos activity between PBS and LPS-conditioned mice on both training and test day (Fig. 3a). We hypothesized that brain regions involved in sickness memory recall would show increased c-Fos activity both in response to LPS on training day and during memory retrieval on test day. To map neuronal activity across multiple brain regions, we used an automated c-Fos detection pipeline (Fig. 3b,c)^35^. On training day, LPS-injected mice showed decreased c-Fos expression in the dentate gyrus (DG), cornu ammonis 3 (CA3), and cornu ammonis 1 (CA1) compared to PBS controls (Fig. 3d-f). On test day, DG activity was similar between groups, reflecting its crucial role in encoding contextual information in general (Fig. 3d). However, we observed elevated c-Fos expression in CA3 and CA1 in LPS-conditioned mice, providing evidence of plasticity within memory-associated circuits (Fig. 3e,f).

**Figure 3:**
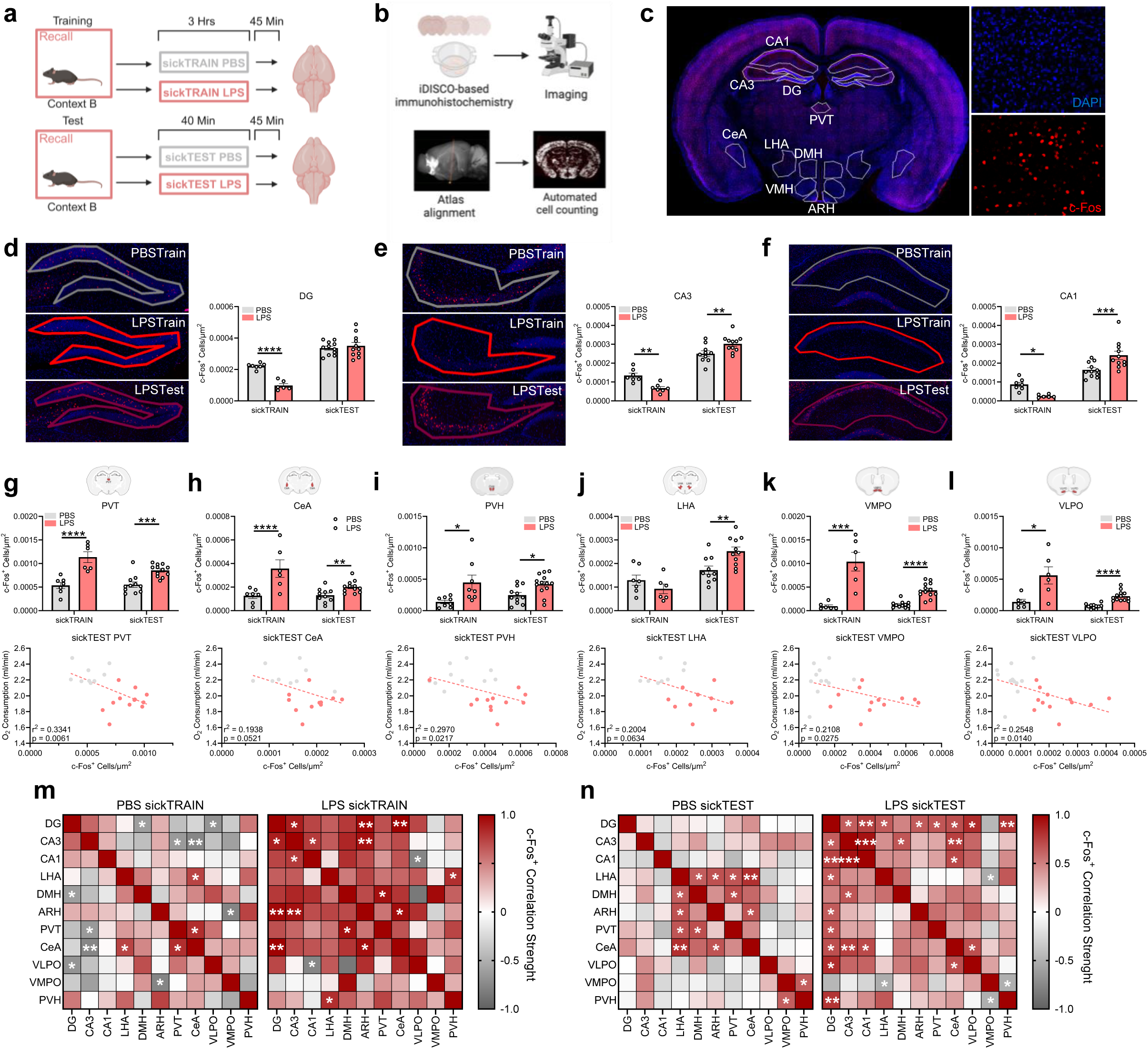
A contextual sickness-memory increases activity in sickness-associated brain regions. **a**, Experimental timeline for tissue collection. **b**, Automated brain-wide c-Fos detection pipeline. **c**, Hippocampal slice (left) with c-Fos^+^ cells (red; bottom right) and DAPI^+^ cells (blue; top right). **d-f**, c-Fos^+^ quantification normalized to region area in the **d**, DG, **e**, CA3, and **f**, CA1 during sickTRAIN and sickTEST between PBS-(grey) and LPS-injected (red) mice (right), with representative regional images (left). **g-l**, c-Fos^+^ quantification normalized to region area in the **g**, PVT, **h**, CeA, **i**, PVH, **j**, LHA, **k**, VMPO, and **l**, VLPO during sickTRAIN and sickTEST between PBS-(grey) and LPS-injected (red) mice (top), correlated with oxygen consumption on test day (bottom). **m**, Correlation matrix of c-Fos^+^ cells normalized to area during sickness training day (sickTRAIN) in PBS-(left) and LPS-injected (right) mice, with positive (red) and negative (grey) correlations. **n**, Correlation matrix of c-Fos^+^ cells normalized to area during sickness recall day (sickTEST) in PBS-(left) and LPS-paired (right) mice, with positive (red) and negative (grey) correlations. Data are shown as mean ± s.e.m, n = 6–12 mice per group. Significance was calculated using a two-way ANOVA (**d-l**), or a simple linear regression (**g-n**). \**P* < 0.05, \*\**P* < 0.01, \*\*\**P* < 0.001, \*\*\*\**P* < 0.0001.

Many brain regions have been shown to display increased activity in response to inflammatory signals during sickness and disease, including the CeA^36,37^, PVT^12^, and PVH^30,38^. Among these, the hypothalamus shows the most robust activation following LPS administration, with areas such as the VMPO regulating fever, body temperature, and appetite via connections with other hypothalamic nuclei, including the arcuate nucleus (ARH)^11,30^. To assess whether sickness-associated regions are engaged during memory encoding and recall, we quantified c-Fos expression on both training and test days. On training day, LPS injection led to elevated c-Fos activity in the PVT, CeA, PVH, lateral hypothalamic area (LHA), VMPO, ventrolateral preoptic nucleus (VLPO), ARH, and dorsomedial hypothalamus (DMH; Fig. 3g-l, top; Extended Data Fig. 4a,b, left). Strikingly, these same regions were reactivated during memory recall in LPS-conditioned mice compared to PBS controls, suggesting plasticity-induced engagement of sickness-relevant circuits (Fig. 3g-l, top; Extended Data Fig. 4a,b, left).

To explore whether activation of these regions relates to systemic physiology, we correlated c-Fos expression in each region with whole-body metabolic rate on test day. We found a strong inverse correlation between c-Fos activity in the PVT, CeA, PVH, LHA, VMPO, VLPO and DMH, suggesting that increased activity in these regions on test day reduced metabolic rates (Fig. 3g-l, bottom; Extended Data Fig. 4a, right). In contrast, although ARH activity was elevated during recall, it did not correlate with metabolic rate (Extended Data Fig. 4b, right). Finally, to explore whether the hippocampus coactivated with sickness regions, we performed correlation analyses of c-Fos dynamics across brain regions. We found strong positive correlations between hippocampal DG, CA1 and CA3 with sickness regions such as the CeA on training day in LPS-administered mice. Similarly, coactivation between the hippocampus and hypothalamic sickness regions increased in the LPS-conditioned mice compared to the PBS conditioned (Figure 3m,n). Additionally, hippocampal activity did not coactivate with just any hypothalamic region (Extended Data Fig. 4c,d). These findings suggest that brain regions activated during acute sickness are re-engaged during contextual sickness memory recall and may contribute to modulation of whole-body metabolism.

### Sickness memory engrams are found in the hippocampus, thalamus and hypothalamus

Previous studies have described effector brain regions which control inflammation and sickness behaviour^10,30^. However, it remains unknown whether inflammatory information can be stored as memory engrams for later retrieval. To address this, we crossed TRAP2 mice with R26R-STOP-floxed enhanced yellow fluorescent protein (eYFP) mice, enabling visualization of sickness engram encoding and retrieval in the hippocampus and hypothalamus (Fig. 4a)^39,40^. To visualise sickness engrams across brain regions, mice underwent our Pavlovian conditioning timeline with 4-OHT delivered on training day in context B (Fig. 4b,c). On test day, mice were either kept in their home cage or returned to context B to trigger memory retrieval. We observed increased neuronal activity in the CeA, LHA, PVT, PVH and VLPO of retrieval mice compared to home cage controls, with no differences in DG activity, consistent with previous results (Fig. 3g-l; Fig. 4f-k, top). Although there was a significant decrease in c-Fos^+^ cells in the DG, CA3 and CA1 on training day, the number of eYFP-labelled cells did not differ between groups. This confirmed equal labelling during the sickness training day, since both groups underwent the same experience (Fig. 4f-k, middle). Finally, we quantified engram reactivation by identifying neurons positive for both eYFP (encoding) and c-Fos (retrieval). Mice recalling the sickness context exhibited significantly greater overlap of eYFP and c-Fos expression in DG, CeA, LHA, PVT, PVH, and VLPO than home cage controls (Fig. 4f-k, bottom). Additionally, we identified sickness memory ensembles in the DMH and ARH (Extended Data Fig. 5a,b). Together, these data indicate that sickness engrams form across the hippocampus, thalamus, and hypothalamus.

**Figure 4:**
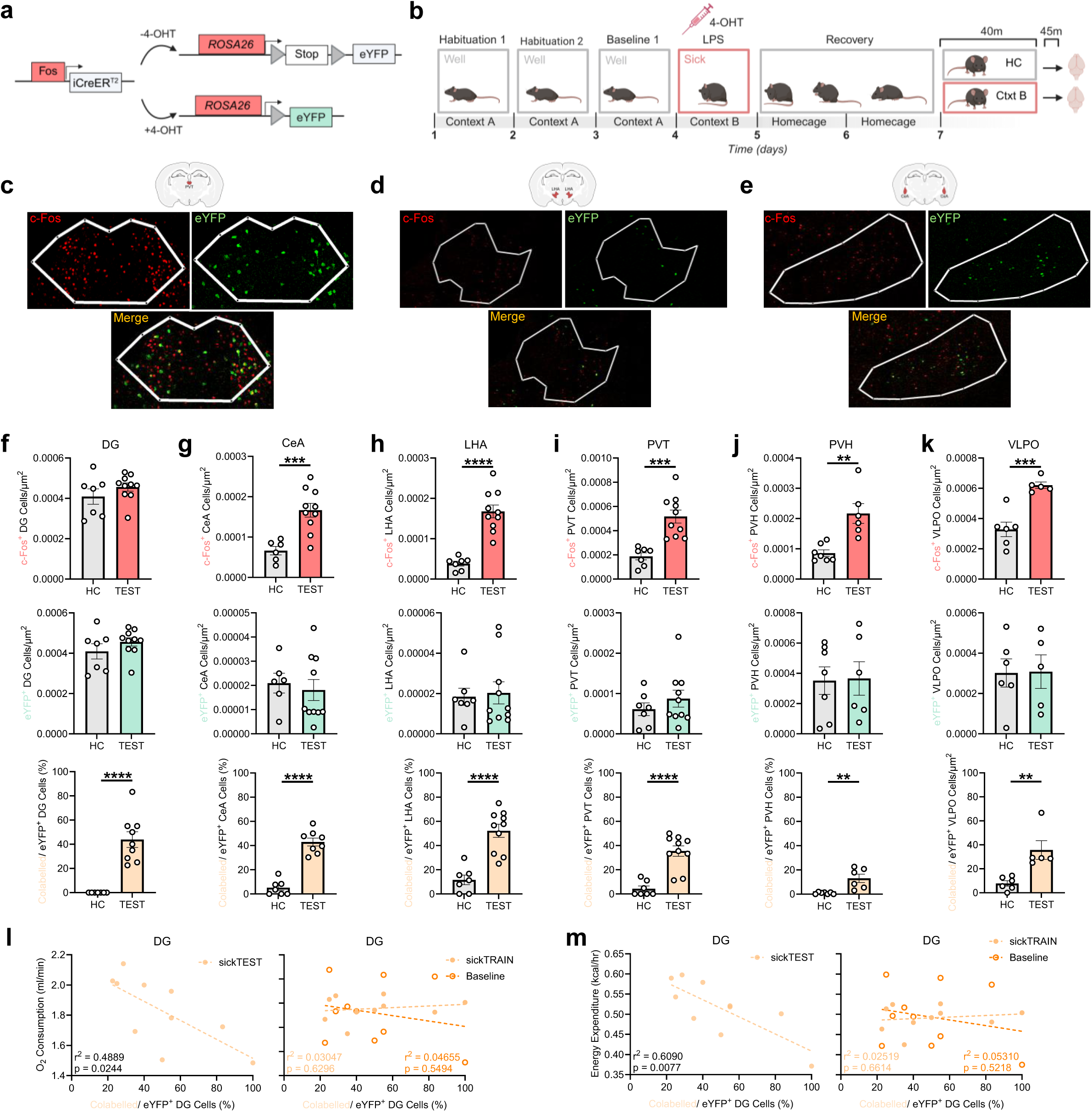
Contextual sickness engrams are in the DG, and sickness-associated brain regions. **a**, Genetic strategy for *Trap2:R26R* mice. **b**, Experimental timeline for engram labelling. **c-e**, Representative images of c-Fos^+^ (red, top-left) cells, eYFP^+^ (green; top-right), and colabelled (yellow, bottom) cells in the PVT (**c**), LHA (**d**) and CeA (**e**). **f-k**, c-Fos^+^ neurons (top), eYFP^+^ neurons (middle), and colabelled neurons (bottom) normalized to region area in the **f**, DG, **g**, CeA, **h**, LHA, **i**, PVT, **j**, PVH and **k**, VLPO between home cage controls (grey) and test (red, green, orange) mice during sickness recall. **l**, Correlation between DG colabelled cells and oxygen consumption on recall day (sickTEST; left), and oxygen consumption on baseline and sickness training (sickTRAIN) days (right). **m**, Correlation between DG colabelled cells and energy expenditure on recall day (sickTEST; left), and oxygen consumption on baseline and sickness training (sickTRAIN) days (right). Data are shown as mean ± s.e.m, n = 5–10 mice per group. Significance was calculated using a two-tailed unpaired students t-test (**f-k**), or a simple linear regression (**l,m**). \*\**P* < 0.01, \*\*\**P* < 0.001, \*\*\*\**P* < 0.0001.

To determine whether engram reactivation influences whole-body metabolic rate on test day, we correlated the degree of engram overlap with oxygen consumption, carbon dioxide emission, and energy expenditure. We found a strong inverse correlation specifically in the dentate gyrus (DG), where greater engram overlap was associated with lower oxygen consumption, carbon dioxide emission, and energy expenditure on test day (Fig. 4I,m; Extended Data Fig. 5c). In contrast, no correlation was observed between sickness engram reactivation and whole-body metabolism on either baseline or sickness training days (Fig. 4I,m; Extended Data Fig. 5c). These findings suggest that reactivation of sickness memory engrams in the DG modulates whole-body metabolic activity, potentially via downstream sickness effector regions such as PVT and CeA.

### Artificial sickness engram activation decreases whole-body metabolism and modulates liver APPs

We next investigated whether sickness engrams in the DG are functionally relevant for whole-body metabolism and a sickness response. To allow for optogenetic manipulation, we labelled PBS and LPS-paired engrams in the DG with channelrhodopsin-2 (ChR2; Fig. 5a). Mice were trained in our sickness paradigm, but were allowed to recover for one week post-LPS to ensure optimal ChR2 expression in DG neurons, before reactivation of engrams in neutral context (Fig. 5b,c; Extended Data Fig. 6a). Optogenetic activation of PBS-paired engrams did not significantly alter oxygen consumption, carbon dioxide emission, or energy expenditure (Fig. 5d; Extended Data Fig. 6b,d). In contrast, reactivation of LPS-paired engrams led to a significant reduction in whole-body metabolism (Fig. 5e; Extended Data Fig. 6c,e). We next asked whether artificial reactivation of sickness engrams modulates activity in downstream sickness effector regions (Fig. 5f). Activation of LPS-paired DG engrams increased activity in the PVT, CeA and LHA compared to PBS-paired controls (Fig. 5g). Together, these data demonstrate that DG sickness engrams are functionally relevant, modulating activity in sickness effector brain regions, and leading to a decrease in whole-body metabolism.

**Figure 5:**
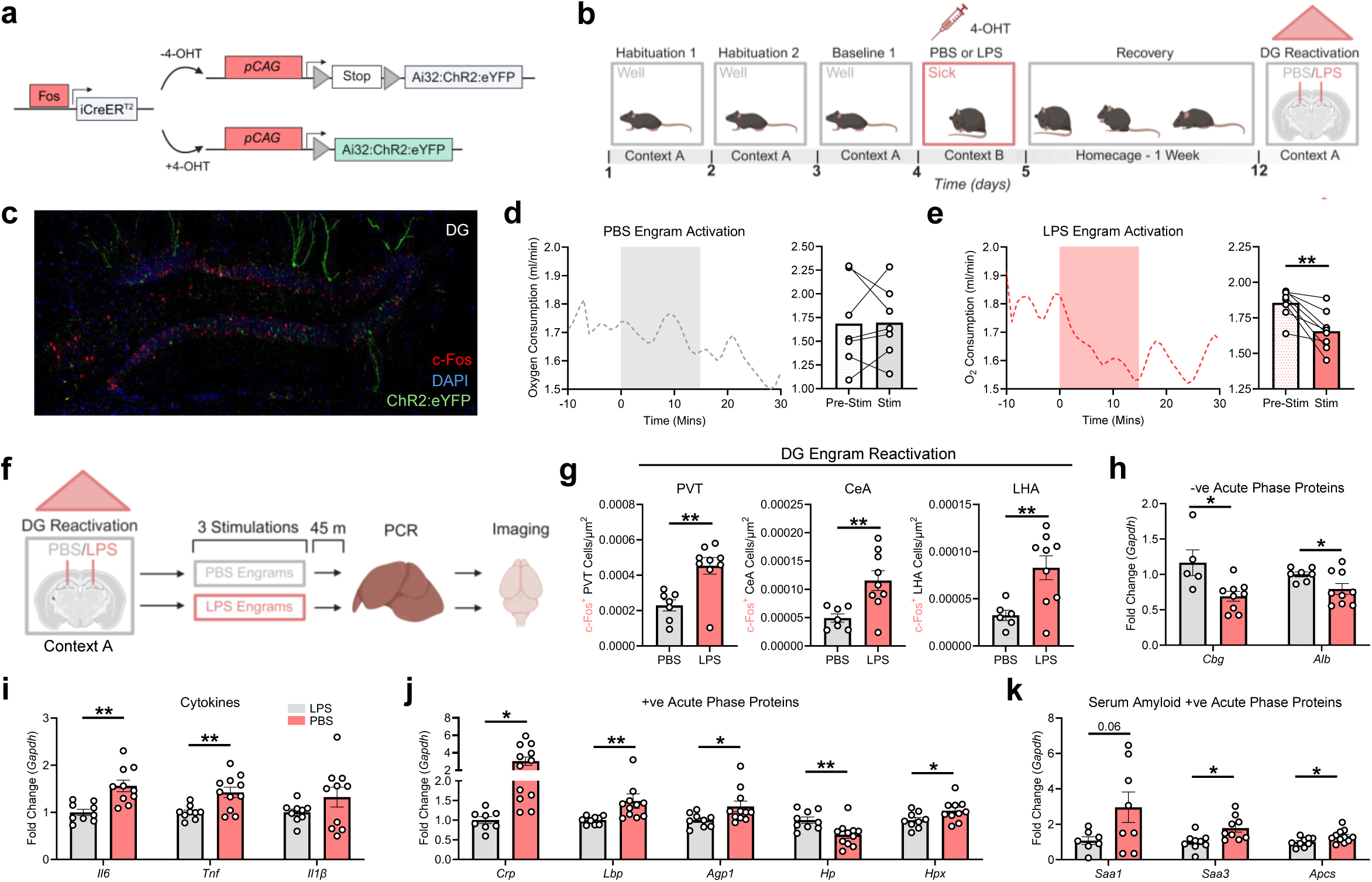
Optogenetic activation of a sickness engram decreases whole-body metabolism and modulates liver APPs. **a**, Genetic strategy for *Trap2:Ai32* mice. **b**, Experimental timeline for optogenetic reactivation (red triangle indicates light activation). **c**, Representative image of hippocampal slice with c-Fos^+^ cells (red) and eYFP^+^ cells (green). **d**, Time plot of oxygen consumption during artificial reactivation of PBS-associated engrams (left), with time averaged pre-light stimulation and during light stimulation (right). Grey overlay indicates laser on. **e**, Time plot of oxygen consumption during artificial reactivation of LPS-associated engrams (left), with time averaged pre-light stimulation and during light stimulation (right). Red overlay indicates laser on. **f**, Experimental timeline of tissue collection. **g**, Comparison of c-Fos^+^ neurons normalized by area in the PVT, CeA and LHA after optogenetic reactivation of PBS-associated (grey) or LPS-associated (red) engrams in the DG. **h,i,** Relative expression of the negative APPs *Cbg* and *Alb* (**h**), and the cytokines *Il6*, *Tnf* and *Il1β* in the liver after reactivation of PBS-associated (grey) or LPS-associated (red) engrams. **j,k,** Relative expression of the positive APPs *Crp*, *Lbp*, *Agp1*, *Hp* and *Hpx* (**j**), and the serum amyloid positive APPs *Saa1*, *Saa3*, and *Apcs* (**k**) in the liver after reactivation of PBS-associated (grey) or LPS-associated (red) engrams. Data are shown as mean ± s.e.m, n = 5– 12 mice per group. Significance was calculated using a two-tailed paired students t-test (**d,e**), or a two-tailed unpaired students t-test (**g-k**). \**P* < 0.05, \*\**P* < 0.01.

Since artificial reactivation of sickness engrams decreased whole-body metabolic rate in LPS-Paired mice, we next asked whether optogenetic activation also modulates APP expression in the liver. Both PBS and LPS-paired mice received three 15-minute stimulations, and livers were collected 45 minutes after the final stimulation (Fig. 5f). Reactivation of LPS-paired engrams in the DG led to a decrease in negative APPs *Cbg* and *Alb*, and an increase in the pro-inflammatory cytokines *Il6* and *Tnf*. Positive APPs, including *Crp*, *Lbp*, *Agp1*, *Hpx*, *Saa1*, *Saa3*, and *Apcs*, were also upregulated compared to PBS-paired controls (Fig. 5h–k). To confirm that these changes were not due to residual effects of prior LPS exposure, a separate cohort of mice was injected with PBS or LPS and allowed to recover for one week before liver tissue was collected (Extended Data Fig. 7a). No significant differences in cytokine, positive APP, or negative APP expression were observed between groups at this time point, suggesting that optogenetic reactivation of sickness engrams is sufficient to modulate liver APP expression (Extended Data Fig. 7b-e).

### Inhibition of sickness engrams prevents a decrease in metabolic rate

Considering that reactivation of sickness engrams decreased whole-body metabolism and modulated liver APP expression, we next examined whether inhibition of sickness engrams could prevent the physiological response to sickness recall. To test this, we administered the inhibitory DREADD (designer receptors exclusively activated by designer drugs) hM4Di into the DG of the hippocampus (Fig. 6a). DREADD expression was targeted to DG neurons active during LPS exposure. On test day, mice received either clozapine-N-oxide (CNO) or saline 30 minutes before being re-exposed to the LPS-paired context B (Fig. 6b,c). Chemogenetic inhibition of DG sickness engrams with CNO prevented the LPS-associated decrease in oxygen consumption, carbon dioxide emission, and energy expenditure compared to saline controls (Fig. 6d-f). Moreover, inhibition of DG sickness engrams prevented activation of sickness associated effector regions compared to saline controls (Fig. 6g). Finally, CNO inhibition of LPS-paired engrams prevented an increase in *Il6* expression in the liver (Fig. 6h). These findings demonstrate the necessity of a contextual sickness engram in the DG to modulate metabolism, sickness region activity, and liver APPs in response to an LPS-paired memory.

**Figure 6:**
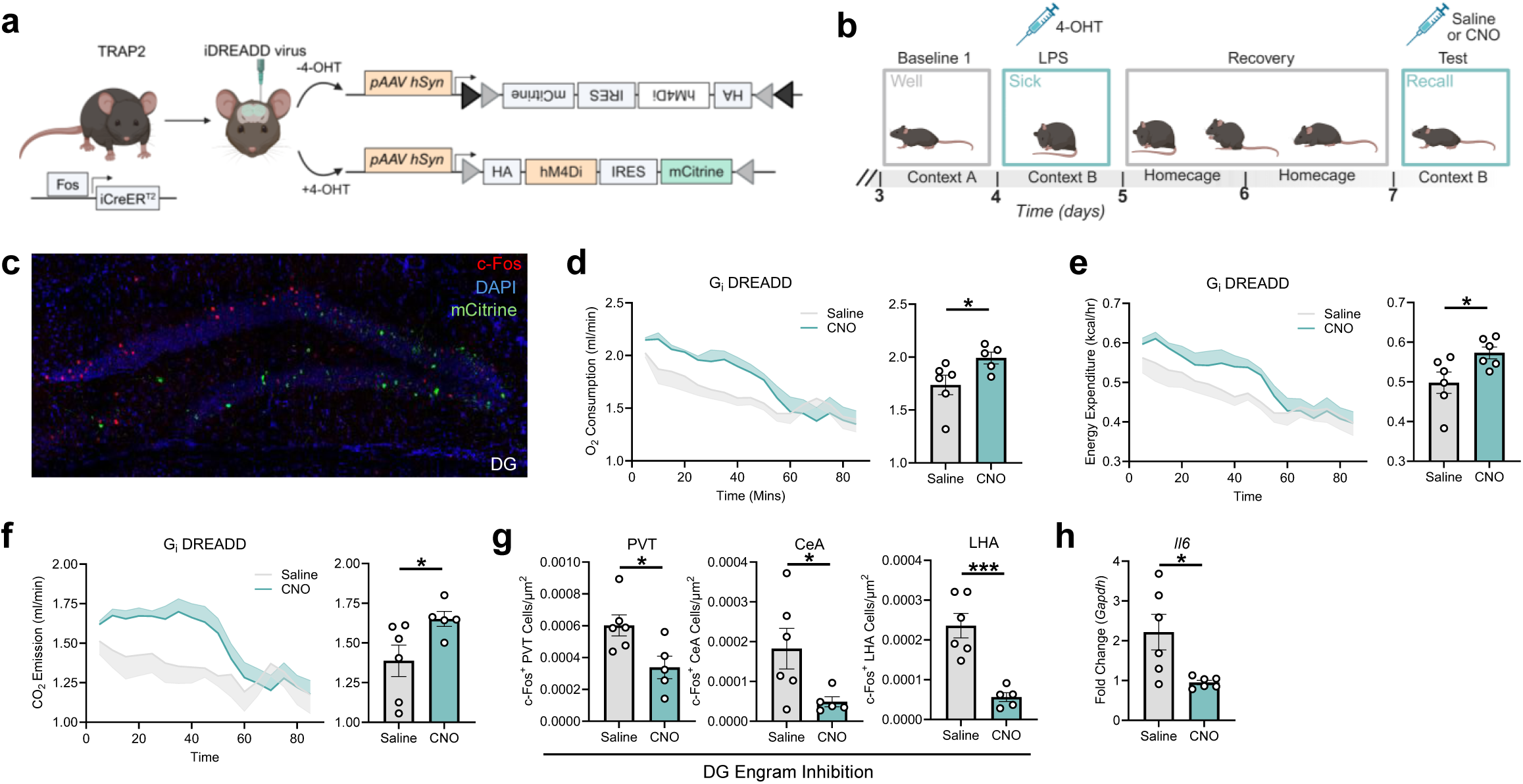
DREADD inhibition of a sickness engram prevents a decrease whole-body metabolism. **a**, Genetic strategy for *Trap2* mice with DREADD injection. **b**, Experimental timeline for DREADD labelling. **c**, Representative image of hippocampal slice with c-Fos^+^ cells (red) and mCitrine^+^ cells (green). **d**, Time plot of oxygen consumption during DREADD inhibition of sickness engrams (left), with averaged time between saline injected (grey) and CNO injected mice (green). **e**, Time plot of energy expenditure during DREADD inhibition of sickness engrams (left), with averaged time between saline injected (grey) and CNO injected mice (green). **f**, Time plot of Carbon dioxide emission during DREADD inhibition of sickness engrams (left), with averaged time between saline injected (grey) and CNO injected mice (green). **g**, Comparison of c-Fos^+^ neurons normalized by area in the PVT, CeA and LHA after DREADD inhibition of sickness engrams in the DG between saline injected (grey) and CNO injected mice (green). **h,** Relative expression of the cytokine *Il6* in the liver after DREADD inhibition of sickness engrams in the DG between saline injected (grey) and CNO injected mice (green). Data are shown as mean ± s.e.m, n = 5–6 mice per group. Significance was calculated using a two-tailed unpaired students t-test (**d-h**). \**P* < 0.05, \*\*\**P* < 0.001.

## Discussion

To survive in the wild, animals must perceive and predict uncertain environments. Sickness events can subsequently lead to decreased food intake, energy, and mobility, thereby increasing the risk of predation. Thus, encoding information about stimuli associated with prior sickness, such as specific food substances, locations or other conspecifics is highly adaptive. In this study, we demonstrate that an LPS injection can be paired with contextual cues such that, after recovery, mice exhibit memory recall of the sickness event when returned to the same context. We show that an LPS-paired conditioned response decreases metabolic rate upon reexposure to the sickness-associated context. The concept of immune conditioning emerged in 1886 with the first description of the Rose-Cold study^14,15^. Since then, immune conditioning has been described using various combinations of conditioned stimuli (contextual information) with unconditioned stimuli (immune modulators)^19–21^. For example, pairing the immunosuppressive agent cyclophosphamide with saccharin-flavored water was sufficient to suppress immune responses upon re-exposure to saccharin alone^41^. Similarly, pairing of LPS with saccharin water could condition a febrile fever response^42^. Our findings extend this body of work by demonstrating, for the first time, that contextual cues can form Pavlovian associations with sickness experiences, resulting in the formation of context-specific sickness engrams that control changes in whole-body metabolism. To our knowledge, this is the first study to use metabolic readouts to assess recall of anticipatory immunity.

Brain-body interactions have become a central topic of recent research, providing fundamental information about sensory inputs feeding the brain information, and the brain directing peripheral responses. In particular, it has been shown that psychological stress modulates immune cell numbers in the blood through activity of the PVH in the brain^43^. More recently, we identified a link between memory and physiological changes where reactivation of cold-engrams in the DG modulates the expression of thermoregulatory genes in brown adipose tissue, enhancing adaptability to cold exposure^35^. In this study, we show that recalling a contextual memory associated with prior sickness modulates the expression of APPs in the liver. To our knowledge, this is the first demonstration of immune conditioning of liver APPs. We hypothesize that the recall of a sickness-associated memory triggers an anticipatory modulation of hepatic APPs, preparing the host for potential secondary immune challenge. However, dissociating the contribution of immunological memory from memory recall in driving anticipatory immunity presents a key conceptual challenge for future work.

There is an ever-increasing body of evidence that the immune system’s response to insults is coordinated by activity in specific brain regions. Recent studies have identified regions in the brainstem^8,10^, hypothalamus^11,30^ and cortex^12,22^ which modulate an immune response. Structures such as the NTS, VMPO and IC differentially regulate aspects of host defence. Activity in NTS neurons regulates the production of pro- and anti-inflammatory cytokines and modulates sickness-associated behaviours^10,30^, while VMPO neurons regulate body temperature^11^. The response of neurons in the NTS and VMPO to LPS demonstrate their role as sickness effector regions. Prior work has demonstrated that neuronal ensembles in the IC are involved in immune responses related to taste aversion and gut inflammation^12,22^. However, whether memory engrams play a direct role in regulating predictive immune function remained unknown. Here, we show for the first time that contextual sickness memories are encoded in the hippocampal DG during LPS exposure, and are reactivated upon memory recall in the absence of LPS. Furthermore, sickness engrams are found beyond the hippocampus in traditional sickness regions such as the PVT, CeA and PVH. Upon optogenetic reactivation, DG sickness engrams modulate whole-body metabolic responses and APP expression in the liver, effects which are reversed by chemogenetic DREADD inhibition. Notably, optogenetic activation and DREADD inhibition increased and decreased c-Fos activity in sickness regions respectively, suggesting an emergent connection between the hippocampus and sickness effector regions after learning.

A key question arising from our findings is whether sickness-related memory traces contribute to enhanced immune responses during subsequent infections or immune challenges. Most likely, anticipatory immunity bolstered by sickness memory would be beneficial for resolving subsequent infections. Additionally, sickness memories may also drive behavioral avoidance of previously associated cues, while simultaneously priming physiological immune responses as a biological safeguard. Supporting this, a recent study demonstrated that the sight of a sick person alone activates networks in motor and salience regions of the human brain, to pre-emptively activate innate lymphoid cells in the periphery^44^. Furthermore, conditioned immune responses have also shown efficacy in preclinical models, where learned immunosuppression prolonged heart transplant engraftment^45^, and attenuated tumour progression^46,47^. Conditioned immune responses have been suggested as the mechanism by which the placebo effect occurs in patients, and may be leveraged as a potential clinical application in conjunction with pharmacological interventions to reduce drug dosages and unwanted side effects. A clinical study implemented learned immunosuppression in kidney transplant patients, where the immune conditioned group displayed decreased T cell proliferation and cytokine production compared to typical treatments^48^. Together these studies show a strong capacity for learned immune responses to be used clinically to treat a host of human diseases ranging from cancer to autoimmunity, which may be modulated by sickness engram activity in humans.

## Acknowledgements

We thank Clara Ortega-de San Luis, Tamara Boto and past and present members of the Ryan Lab for scientific discussions and support. We thank Bartosz Schramm, Gavin McManus, Barry Moran and Rustam Rakhmatullin for their continuous technical support.

## Author Contributions

Conceptualization: A.D., A.M.Z., T.J.R. Experimental design: A.D., A.M.Z., P.C., L.L., T.J.R. Methodology: A.D., A.M.Z., P.C., L.L., T.J.R. Investigation and analyses: A.D., A.M.Z., P.C, E.U. Data interpretation: A.D., A.M.Z., L.L., T.J.R. Funding acquisition: T.J.R. Project administration: A.D., T.J.R. Supervision: L.L., T.J.R. Writing of the original draft: A.D., T.J.R. All authors contributed to writing and editing of the final draft.

## Inclusion and Diversity

We support inclusive, diverse, and equitable conduct of research.

## Methods

### Mice

In all experiments, male C57BL/6J-OlaHsd mice between 8 and 12 weeks of age were used. Experiments involving engram labelling were performed using *Trap2:Ai32*^39,49^ and *Trap2:R26R*^40,50^ transgenic mice. Mice were group housed at 22 °C under a 12-hour light–dark cycle (lights on 07:00– 19:00), with *ad libitum* access to food and water. All testing was performed during the light phase. Mice were bred in-house under specific-pathogen-free conditions in accordance with Irish and European Union regulations. All mouse work was performed in compliance with the T.R. laboratory project licence, with ethical approval from the Trinity College Dublin ethics committee and the Animal Research Ethics Committee from the Health Products Regulatory Authority (HPRA). For experiments using lipopolysaccharide (LPS; Merck), mice were administered a single 500μg/kg dose, or PBS control, by intraperitoneal injection.

### Conditioned place aversion

The conditioned place aversion apparatus consisted of two separate chambers with specific contextual cues. Mice could freely explore each side through an opening in the centre wall between the two chambers. On day 1, mice were placed inside the apparatus to freely explore both sides for a total of 20 min. On day 2, the centre opening of the apparatus was closed, and mice were injected with either PBS or LPS just prior to being placed into one side of the apparatus for 4 hours, such that the mice were getting gradually sick in the context. Mice were then returned to their home cage for 2 days to recover from the LPS-induced sickness. On day 5, the centre wall was re-opened and mice were placed back into the middle apparatus to freely explore both chambers for 20 minutes. The behaviour on both day 1 (pre-test) and day 5 (test) were videotaped and later hand scored for the time spent in each chamber. The time spent in each chamber was normalized using the formula to account for any pre-existing preferences (test phase duration spent in the cold-paired chamber/time spent there in the pre-test phase). Each side of apparatus was counterbalanced across all mice for both PBS and LPS groups.

### Sickness training schedule

Male mice were individually housed in the Promethion cages for 6-hours per day at 21°C. For habituation to the metabolic cages, mice were placed in the Promethion cages on days 1 and 2. On day 3, mice were put into the Promethion cages while metabolic and behavioural data were simultaneously recorded to establish baseline measurements (context A). On day 4, contextual cues were added to the chambers consisting of wall and floor patterns, brighter lights, different bedding and an acetic acid odour (context B). Just prior to being placed into context B, mice were injected with either PBS or LPS to pair context B with sickness. The mice spent a total of approximately 6-hours in context B, with metabolic and behavioural data recorded throughout. Mice were then returned to their home cages for a two-day recovery period. On day 9, mice were placed back into context B inside the metabolic cages after recovery from sickness, with metabolic and behavioural data simultaneously recorded to establish the test-day measurements.

### Metabolic cages

Indirect calorimetry experiments were performed on singly housed mice in Promethion metabolic cages (Sable Systems). O_2_ consumption, CO_2_ emission, energy expenditure, body weight, food and water intake and locomotor activity were monitored throughout the 8-hour session each day. Mice had *ad libitum* access to food and water throughout the entire session. Respiratory gasses were measured with an integrated fuel cell oxygen analyser, a spectrophotometric carbon dioxide analyser and a water vapour pressure analyser (GA3m1, Sable Systems). Before each run, gas sensors were calibrated with 100% N_2_ as zero reference and with a span gas containing a known concentration of 0.933% CO_2_. Air flow was controlled with the multichannel mass flow generator (FR8-1, Sable Systems). Flow rate was set at 2,000 ml min^−1^ for each individual cage. Oxygen consumption and carbon dioxide production were measured for each mouse at 3–4-min intervals. Energy expenditure was calculated using the equation: K_cal/h_ = 60 x (0.0003941 x VO_2_ + 0.001106 x VCO_2_)^51^.

### Tissue collection and preparation

Brains and a piece of liver were collected from mice that through the behavioural sickness timeline as described above. Brains and liver were collected from PBS and LPS administered mice on sickness training day (sickTRAIN; day 4) and test day (sickTEST). Brains and liver were collected 3 hour and 45 minutes after PBS and LPS injection on sickness training day, and after 1 hour and 25 minutes on test day. These timings were selected to get an accurate representation of real time sickness on sickness training day, and to get a representation of brain activity after the largest difference in metabolism between PBS-paired and LPS-paired mice on test day.

#### Brains

Mice were transcardially perfused with 50mL of phosphate-buffered saline (PBS), followed by 50mL of 4% paraformaldehyde. Brains were removed and placed into 4% paraformaldehyde for overnight fixation. The next day, brains were transferred into PBS and stored at 4°C until sectioning. Brains were subsequently sectioned as 100-μm coronal sections with a vibratome (Leica VT1200 S) into PBS, and stored in PBS at 4°C until immunostaining.

#### Liver

Mice were transcardially perfused with 50mL of PBS, and a piece of liver was taken, immediately snap frozen in liquid nitrogen and stored at −80 °C. Care was taken to ensure the piece was taken from the same lobe of each liver (medial lobe of mouse liver).

### RNA extraction from tissues

Livers were snap-frozen in liquid nitrogen, defrosted at room temperature and transferred to a 2mL tube containing a 5-mm stainless steel bead. Tissues were homogenized in 1mL trizol reagent (Thermo Fisher Scientific) in a tissue lyser for 2.5 min, 25 pulses per second. Then, 200 µl chloroform was added to each tube, tubes were mixed by inverting and left at room temperature for 2–3 min, before centrifuging at 13,500*g* for 15 min. The aqueous phase containing RNA was transferred into a new Eppendorf tube and 500 µl isopropanol was added to precipitate the RNA. Tubes were inverted ten times and left at room temperature for 10 min, and then centrifuged at 13,500*g* for 10 min. Supernatants were discarded and RNA pellets were washed in 1 ml 75% ethanol 3 times, then resuspended in RNAse free dH_2_O. Tubes were centrifuged at 13,500g for 5 min and supernatants were discarded by inverting the tube. The RNA pellet was left to dry at room temperature for 10–20 min, and the pellets were resuspended in 100µL RNAse free water. RNA was left on ice for 30 min, then in a heat block set at 55 °C for 15 min. RNA quality and concentration were determined using a Nanodrop 2000 UV spectrophotometer (Thermo Fisher Scientific). 20µL of cDNA was synthesized from 2µg of isolated RNA using the High-Capacity cDNA Reverse Transcription Kit (Biosciences) in a MiniAmp Thermal Cycler (BD Biosciences). To quantify the relative mRNA expression of genes of interest, quantitative PCR was performed in 384-well plates (Thermo Fisher Scientific) using SYBR Green-based detection (eBioscience). Relative mRNA levels were calculated using the ΔΔ cycle threshold (ΔΔCt) for 5 min and supernatants were discarded by inverting the tube. The RNA pellet was left to dry at room temperature for 10–20 min, and the pellets were resuspended in 100µL RNAse free water. RNA was left on ice for 30 min, then in a heat block set at 55 °C for 15 min. RNA quality and concentration were determined using a Nanodrop 2000 UV spectrophotometer (Thermo Fisher Scientific). 20µL of cDNA was synthesized from 2µg of isolated RNA using the High-Capacity cDNA Reverse Transcription Kit (Biosciences) in a MiniAmp Thermal Cycler (BD Biosciences). To quantify the relative mRNA expression of genes of interest, quantitative PCR was performed in 384-well plates (Thermo Fisher Scientific) using SYBR Green-based detection (eBioscience). Relative mRNA levels were calculated using the ΔΔ cycle threshold (ΔΔCt) method and normalized to corresponding endogenous controls (*Gapdh*). For a list of primer sequences used, see supplementary table 1.

### Immunohistochemistry

For immunolabelling of sectioned brain slices, we used an iDISCO-based immunohistochemistry protocol as previously described^35,52,53^. Sections were washed three times in PBS, 10 minutes per wash, and subsequently dehydrated in 50% MeOH/PBS at room temperature for 2.5 hours. Sections were then washed three times in 0.2% PBS/Triton X-100 (PBST; Merck) and blocked in PBST containing 10% dimethyl sulfoxide (DMSO; Merck) and 6% normal goat serum (NGS) for 2 hours at room temperature. After blocking, slices were washed three times in PBS with 0.2% Tween-20 (Merck) and 10μg/mL heparin (PTwH; Fisher Scientific). Sections were then incubated with primary antibodies (chicken polyclonal anti-GFP (1:1,000; Invitrogen) and rabbit polyclonal IgG anti-FOS (1:1,000, Synaptic Systems)) at 4 °C on a shaker for three days in PTwH with 5% DMSO and 3% NGS. Sections were washed three times in PTwH and incubated with secondary antibodies (Alexa Fluor 568 anti-rabbit IgG (1:500, Invitrogen) and Alexa Fluor 488 anti-chicken IgG (1:500, Invitrogen) overnight at 4 °C in the dark. The next day, sections were washed three times in PTwH, followed by three washes in PBS at room temperature, ten minutes per wash. Finally, sections were stained with DAPI in PBS for 10 minutes followed by a final wash in PBS, and then mounted on superfrost slides (Fisher Scientific) using Vectashield (2Bscientific), cover-slipped and sealed.

### Confocal imaging

All images were taken on a confocal scanning microscope (Leica TCS SP8, Leica Microsystems). Fluorescence from DAPI was detected at 417–488 nm, Alexa Fluor 568 was detected at 595–650 nm and Alexa Fluor 488 was detected at 500–550 nm. Sections were imaged with a dry 20X objective (NA 0.70, working distance 0.5 mm), with a pixel size of 1.14 × 1.14 μm^2^, a *z* step of 3μm and a *z*-stack of approximately 25μm. Fields of view were stitched together to form tiled images using an automated stage and the algorithm of the LAS X software.

### Automated cell counting

To perform automatic brain-wide analysis of FOS^+^ cells, we used the NeuroInfo software (MBF Bioscience). In brief, immunohistochemistry-labelled whole-brain sections were aligned to the Allen Mouse Brain Atlas using the in-program registration tool. Brain sections were matched to the most closely corresponding atlas plate, but manually adjusted where necessary to ensure accurate fit. Anatomical regions of interest were defined and measured for size in μm^2^ to later normalized cell counts (cells per μm^2^). Next, c-Fos^+^ cells were identified using the cell-detection workflow. For size consistency, only cells between 7 and 19μm were counted and combined with a deep-learning algorithm to successfully identify c-Fos^+^ cells. Detected cells were mapped to the Allen Mouse Brain Atlas and tallied to the corresponding brain structure.

### TRAP2 labelling strategy

The TRAP2 system enables the permanent labelling of neurons that were activated by a given experience^54^. TRAP2 relies on an immediate early gene locus to drive the expression of iCre recombinase combined with a transgenic Cre-dependent effector. The iCre recombinase can be controlled by 4-OHT, a tamoxifen analogue. As such, when a neuron is active in the presence of 4-OHT, the iCre recombinase enters the nucleus, resulting in expression of the effector^39^. For optogenetic experiments, TRAP2 mice were bred with the *Ai32* mice^49^. For engram cell-counting experiments, TRAP2 mice were crossed with the *R26R* mice^40^. To label the cells that were activated during sickness in *Trap2:Ai32* and *Trap2:R26R* transgenic mice, mice were intraperitoneally injected with 4-OHT (20mg/kg, Santa Cruz) and LPS at the same time, before being placed into the Promethion cages on sickness training day (Day 4). As such, all cells that were active during the initial 3 hours of sickness, while the mice were getting progressively sicker, were labelled. Mice remained in the metabolic cages for a total of 6 hours, and then returned to their home cages for recovery. 4-OHT was freshly prepared on the labelling day in a mix of sunflower seed oil and castor oil (4:1, Sigma-Aldrich) at 10 mg/mL.

### Engram tagging and counting

For sickness engram tagging we used *Trap2;R26R* mice. *Trap2;R26R* mice were chosen as they display more reliable labelling in the hypothalamus than other transgenic lines that were tested. All mice were intraperitoneally injected with 4-OHT and LPS at the same time, before being placed into the Promethion cages on sickness training day (Day 4). Mice were then allowed to recover from LPS-induced sickness in their home cages for 2 days. On test day, half of the mice were reexposed to context B, and the other half were kept in their home cage. All mice were euthanized 1 hour 25 minutes into test day recordings for c-Fos quantification using Fiji^55^. eYFP^+^, c-Fos^+^ and colabelled cells were counted bilaterally in the DG, LHA, CeA, PVT, PVH, VLPO, VMPO and DMH. All cell counts were normalized to their respective areas and plotted as cells per μm^2^.

### Stereotactic surgeries

All surgeries were performed when the mice were approximately 7–8 weeks old. 30 minutes prior to surgery mice were subcutaneously injected with meloxicam (15mL/kg) Mice were anaesthetized using a vaporiser to generate a 3% isoflurane concentration and head-fixed on a stereotaxic frame. The isoflurane concentration was then lowered to 1.5% for maintenance throughout the remainder of the surgery. Ophthalmic ointment was applied to the eyes to prevent drying, and intracutaneous lidocaine was injected in the skull. The surgical area was washed with chlorohexidine and 70% ethanol three times before using a blade to perform a 1-1.5 cm incision through the midline of the scalp. The exposed skull was swapped with ethanol and hydrogen peroxide to enhance the visualisation of lambda and bregma. Bilateral craniotomies were performed using a 0.5-mm diameter drill at –2.00 mm anteroposterior and ±1.35 mm mediolateral.

For optogenetic experiments mice were implanted with a custom implant containing two optic fibres (200mm core diameter; Doric Lenses). The optic fibre implant was lowered above the injection site at – 1.75mm dorsoventral. To secure the implant, an even layer of Metabond (C&B Metabond) was applied and left to dry for 15 min. A protective cap was then made from a black polypropylene microcentrifuge tube and secured with dental cement. Mice were allowed to recover from the anaesthesia in a heat chamber at 29 °C before being returned to their home cage. Mice were allowed to recover from surgery for approximately 2–3 weeks before behavioural testing.

For chemogenetic experiments involving DREADDs, TRAP2 mice were injected with 300μl pAAV-hSyn-DIO-HA-hM4D(Gi)-IRES-mCitrine on each side using a microsyringe pump (UMP3; WPI) and a Hamilton syringe (701LT; Hamilton) at –2.00 mm anteroposterior, ±1.35 mm mediolateral and –2.00 mm dorsoventral. The injection speed was 60nL/min and the needle was left inside for an additional 10 minutes after virus delivery to achieve maximum virus spread.

### Optogenetic reactivation of sickness engrams

Optogenetic activation was performed through a 450-nm laser diode fibre light source (Doric LDFLS 450/080). Sickness engrams were labelled on sickness training day (Day 4) and optogenetic reactivation was performed one week later in a neutral context. Mice with optic fibre implants were attached to a patch cord (MFP_200/240/900-0.22_0.3m_FC-ZF1.25(F)) for 30-minute intervals across 3 days for habituation on the days prior to optogenetic reactivation. On reactivation day, mice were exposed to an optogenetic stimulation (8 mA) session that was divided into 3-minutes on and 1-minute off periods for a total of 15 minutes. During the light-on period, mice received blue-light stimulation (20 Hz) with a pulse width of 15ms, and no light stimulation during the light off period. Each 15-minute stimulation period was followed by a 1-hour break with no light. Mice received a total of three rounds stimulation while metabolic rates were simultaneously measured.

### Chemogenetic inhibition of sickness engrams

To inhibit sickness engrams, all mice were placed through the sickness training paradigm, with sickness engrams being labelled on sickness training day (Day 4). All mice were returned to their home cages for 2 days of recovery after labelling. On test day, half of the mice were injected with CNO (2mg/kg, Tocris) and half of the mice were injected with saline 30 minutes prior to being reexposed to the sickness context (Context B). Metabolic measurements were recorded for the duration of the test day.

### Statistics

GraphPad Prism 10 was used for statistical analysis. For all experiments, a 95% confidence interval was used and P ≤ 0.05 was considered statistically significant. A D’Agostino–Pearson omnibus normality test was first performed to test whether the data were normally distributed (Gaussian distribution). If data were normally distributed, parametric testing was performed. If data were not normally distributed, non-parametric testing was performed. When comparing two groups, an unpaired or paired two-tailed student’s t-test was used. When comparing more than two groups, an ordinary one-way ANOVA with Dunnett’s test was used. When comparing data with two variables, a two-way ANOVA with Bonferroni test was used. Simple linear regression was used for correlation analysis, for multiple parameters a corrletaion matrix was generate with the computed *r*^2^ value. In all figures, \**P* ≤ 0.05, \*\**P* ≤ 0.01, \*\*\**P* ≤ 0.001, \*\*\*\**P* ≤ 0.0001.

### Graphical Representation

All diagrams throughout the manuscript were created using Biorender.

## Figures Legends

**Extended Data Fig 1:**
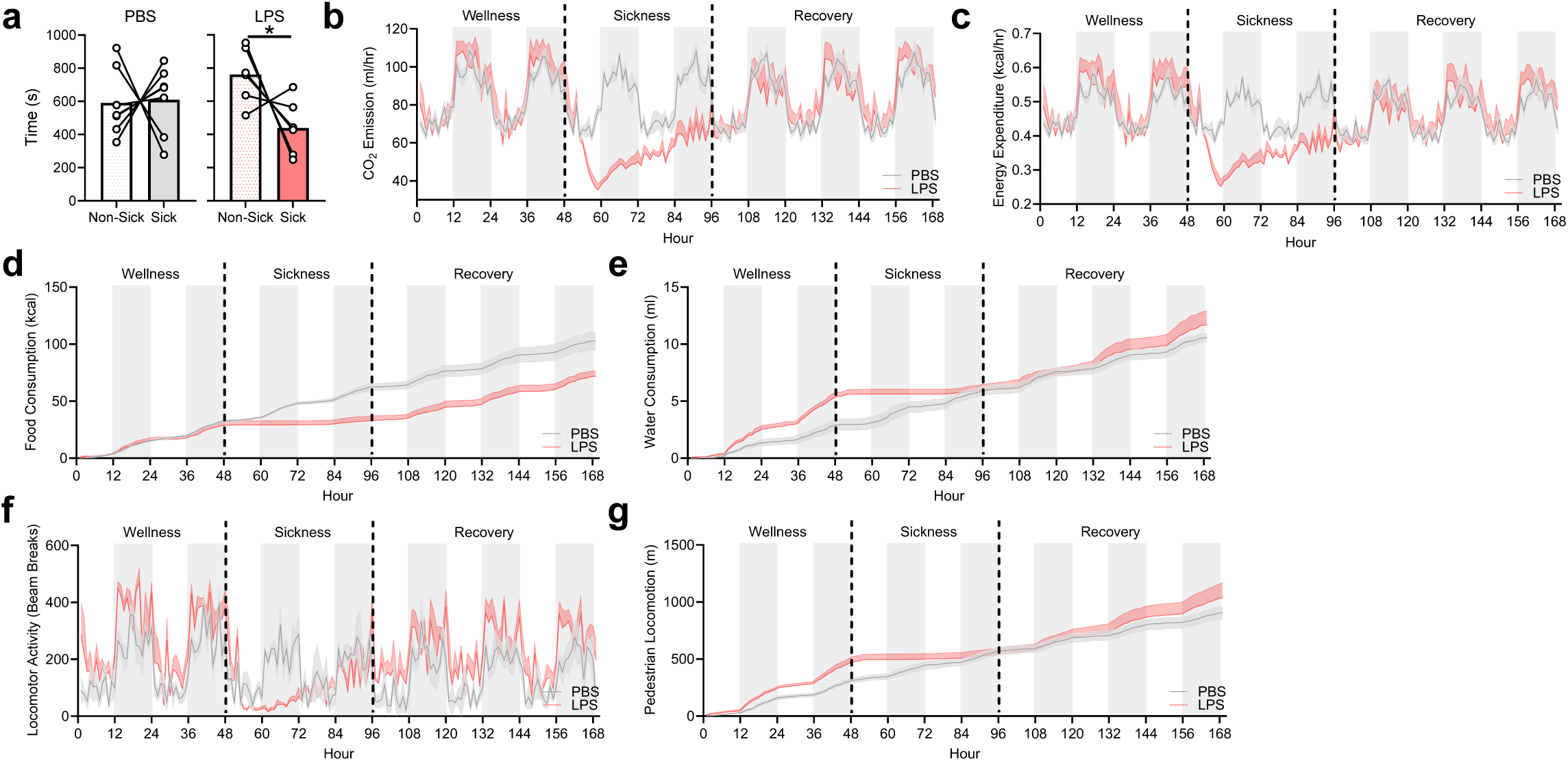
An LPS-mediated immune response decreases whole-body metabolic activity. a, Place preference time for PBS-conditioned aversion (left) and LPS-conditioned aversion (right). b-g, Time plot of (b) carbon dioxide emission, (c) energy expenditure, (d) food consumption, (e) water consumption, (f) locomotor activity, and (g) pedestrian locomotion between PBS-injected mice (grey) and LPS injected mice (red). Data are shown as mean ± s.e.m, n = 4–9 mice per group. Significance was calculated using a two-tailed paired students t-test (a). \**P* < 0.05.

**Extended Data Fig 2:**
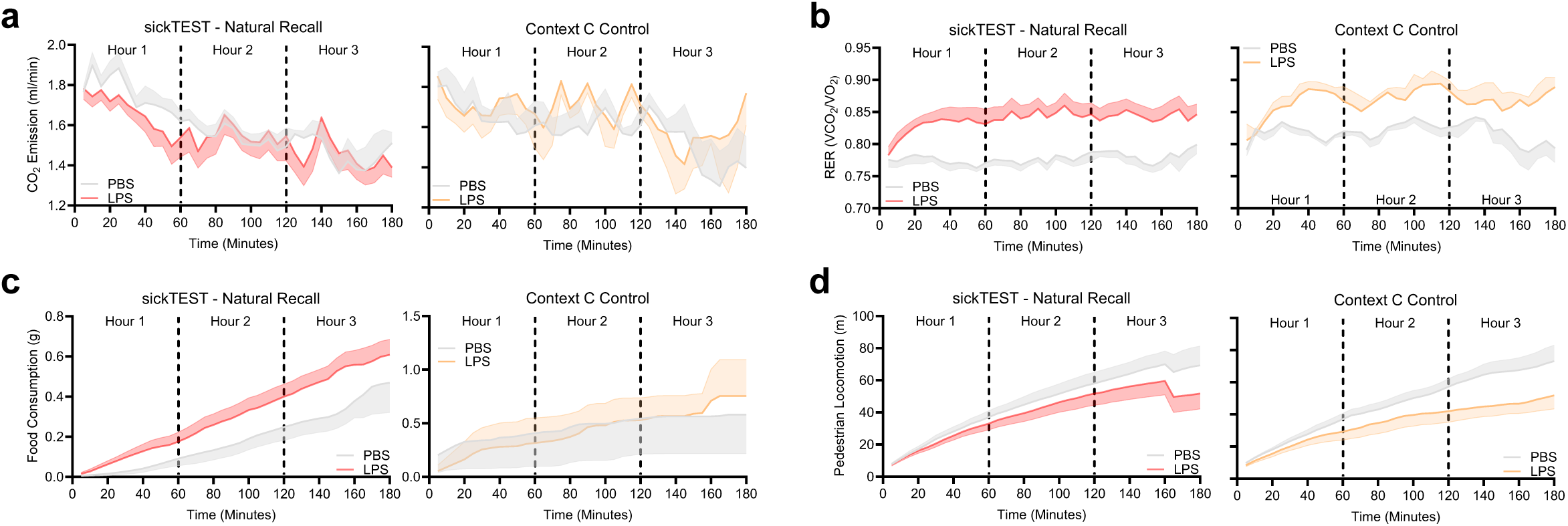
A sickness memory decreases whole-body metabolism in a contextual manner. **a**, Time plot of carbon dioxide emission between PBS-injected mice (grey) and LPS-injected mice (red) on sickness recall day (left), with context C control (right). **b**, Time plot of RER between PBS-injected mice (grey) and LPS-injected mice (red) on sickness recall day (left), with context C control (right). **c**, Time plot of food consumption between PBS-injected mice (grey) and LPS-injected mice (red) on sickness recall day (left), with context C control (right). **d**, Time plot of pedestrian locomotion between PBS-injected mice (grey) and LPS-injected mice (red) on sickness recall day (left), with context C control (right). Data are shown as mean ± s.e.m, n = 4–9 mice per group.

**Extended Data Fig 3:**
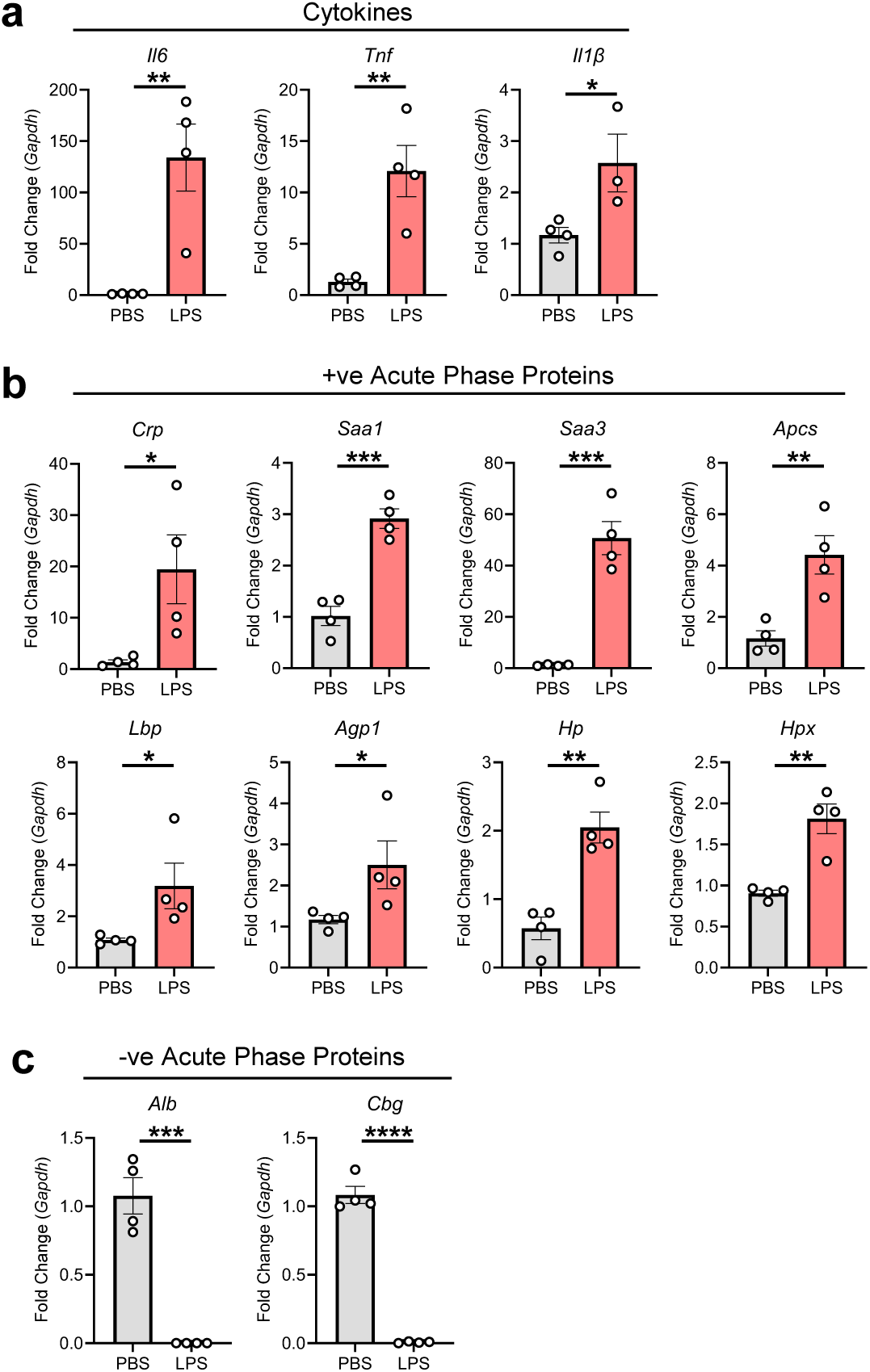
An LPS-mediated immune response modulates liver acute phase response proteins. **a**, Relative expression of the cytokines *Il6*, *Tnf* and *Il1β* in the liver of PBS-(grey) or LPS-injected (red) mice on sickness training day. **b,** Relative expression of the positive APPs *Crp*, *Saa1*, *Saa3*, *Apcs, Lbp*, *Agp1*, *Hp* and *Hpx* in the liver of PBS-(grey) or LPS-injected (red) mice on sickness training day. **c** Relative expression of the negative APPs *Alb* and *Cbg* in the liver of PBS-(grey) or LPS-injected (red) mice on sickness training day. Data are shown as mean ± s.e.m, n = 4 mice per group. Significance was calculated using a two-tailed unpaired students t-test (**a-c**). \**P* < 0.05, \*\**P* < 0.01, \*\*\**P* < 0.001, \*\*\*\**P* < 0.0001.

**Extended Data Fig 4:**
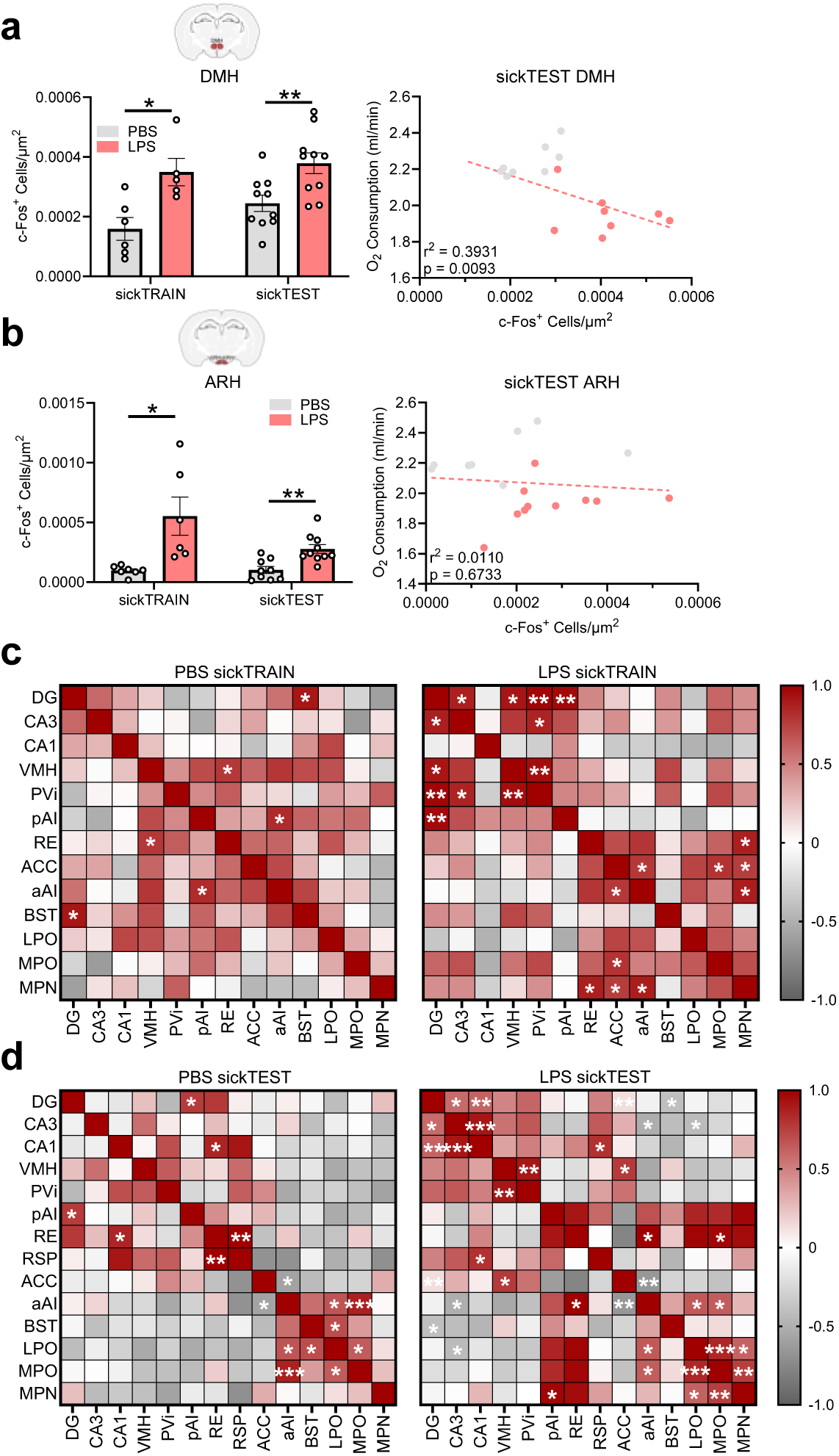
DG activity does not correlate with the activity of just any region on sickTEST. **a**,**b,** c-Fos^+^ quantification normalized to region area in the **a**, DMH and **b**, ARH during sickTRAIN and sickTEST between PBS-(grey) and LPS-injected (red) mice (left), correlated with oxygen consumption on test day (right). **c**, Correlation matrix of c-Fos^+^ cells normalized to area during sickness training day (sickTRAIN) in PBS-(left) and LPS-injected (right) mice, with positive (red) and negative (grey) correlations. **d**, Correlation matrix of c-Fos^+^ cells normalized to area during sickness recall day (sickTEST) in PBS-(left) and LPS-paired (right) mice, with positive (red) and negative (grey) correlations. Data are shown as mean ± s.e.m, n = 5–12 mice per group. Significance was calculated using a two-way ANOVA (**a,b**), or a simple linear regression (**a,b**). \**P* < 0.05, \*\**P* < 0.01.

**Extended Data Fig 5:**
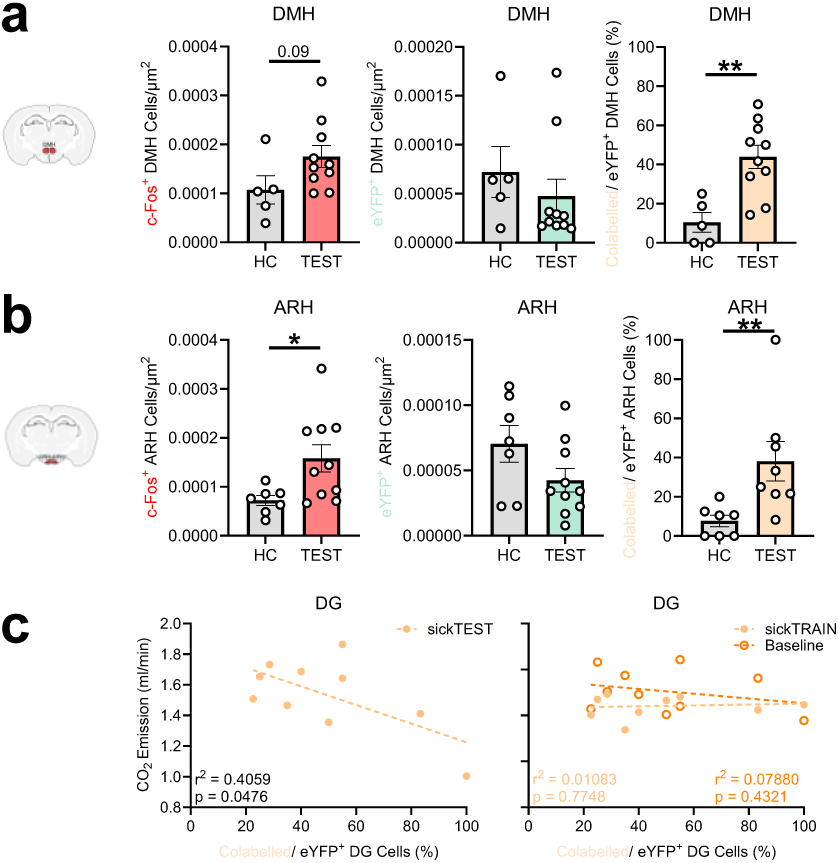
Sickness engrams are in the DMH and ARH of the hypothalamus. **a**,**b,** c-Fos^+^ neurons (left), eYFP^+^ neurons (middle), and colabelled neurons (right) normalized to region area in the **a**, DMH and **b**, ARH between home cage controls (grey) and test (red, green, orange) mice during sickness recall. **c**, Correlation between DG colabelled cells and carbon dioxide emission on recall day (sickTEST; left), and carbon dioxide emission on baseline and sickness training (sickTRAIN) days (right). Data are shown as mean ± s.e.m, n = 5–10 mice per group. Significance was calculated using a two-tailed unpaired students t-test (**a,b**), or a simple linear regression (**c**). \**P* < 0.05, \*\**P* < 0.01.

**Extended Data Fig 6:**
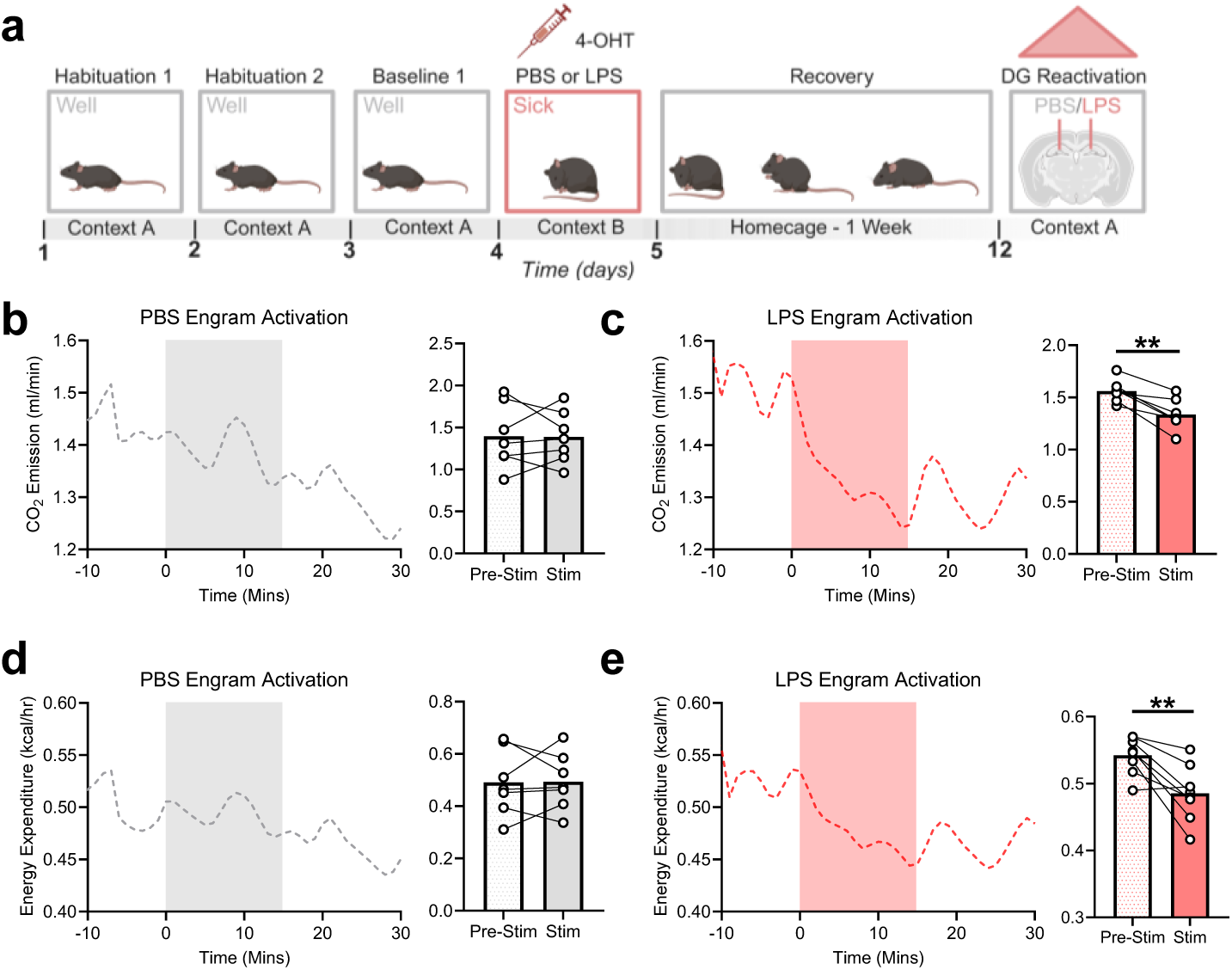
Artificial activation of a sickness engram decreases whole-body metabolism. **a**, Experimental timeline for optogenetic reactivation. **b**, Time plot of carbon dioxide emission during artificial reactivation of PBS-associated engrams (left), with time averaged pre-light stimulation and during light stimulation (right). Grey overlay indicates laser on. **c**, Time plot of carbon dioxide emission during artificial reactivation of LPS-associated engrams (left), with time averaged pre-light stimulation and during light stimulation (right). Red overlay indicates laser on. **d**, Time plot of energy expenditure during artificial reactivation of PBS-associated engrams (left), with time averaged pre-light stimulation and during light stimulation (right). Grey overlay indicates laser on. **e**, Time plot of energy expenditure emission during artificial reactivation of LPS-associated engrams (left), with time averaged pre-light stimulation and during light stimulation (right). Red overlay indicates laser on. Data are shown as mean, n = 8 mice per group. Significance was calculated using a two-tailed paired students t-test (**b-e**). \*\**P* < 0.01.

**Extended Data Fig 7:**
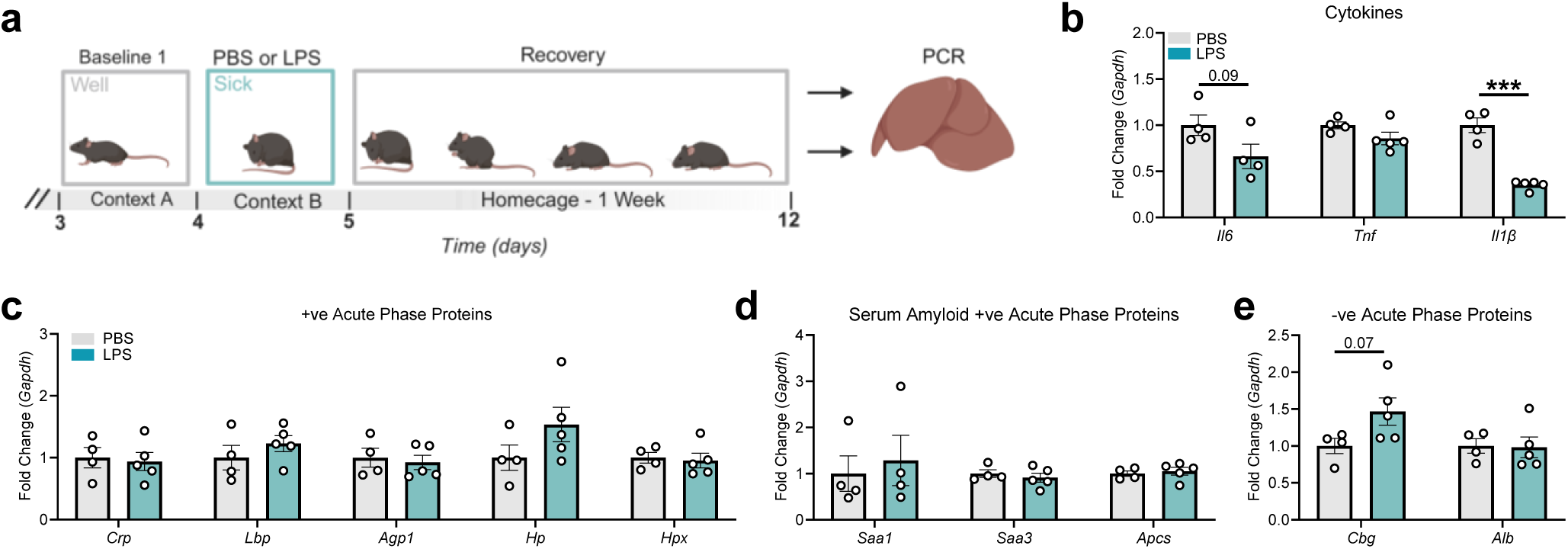
Increased APP gene expression following artificial sickness engram activation is not due to long-term LPS effects. a, Experimental timeline for tissue collection for prolonged effects of LPS. b, Relative expression of the cytokines *Il6*, *Tnf* and *Il1β* in the liver of PBS-(grey) or LPS-injected (green) mice 7 days post-LPS injection. Relative expression of the positive APPs *Crp*, *Lbp*, *Agp1*, *Hp* and *Hpx* (c), the serum amyloid positive APPs *Saa1*, *Saa3*, and *Apcs* (d), and the negative APPs *Cbg* and *Alb* (e) in the liver of PBS-(grey) or LPS-injected (green) mice 7 days post-LPS injection. Data are shown as mean ± s.e.m, n = 4– 5 mice per group. Significance was calculated using a two-tailed unpaired students t-test (b-e). \*\*\**P* < 0.001.

